# Hydrogenase-driven ATP synthesis from air

**DOI:** 10.1101/2025.03.14.643271

**Authors:** Sarah Soom, Stefan Urs Moning, Gregory M. Cook, James P. Lingford, Ashleigh Kropp, Sieu Tran, Rhys Grinter, Chris Greening, Christoph von Ballmoos

## Abstract

All cells require a continuous supply of the universal energy currency, adenosine triphosphate (ATP), to drive countless cellular reactions. The universally conserved F_1_F_o_-ATP synthase regenerates ATP from ADP and P_i_ by harnessing a transmembrane electrochemical proton gradient (*pmf*). Bacteria have evolved a wide diversity of *pmf*-forming proteins to make ATP using light, organic, and inorganic energy sources. Recently, we proposed that many bacteria survive using atmospheric trace gases to produce ATP when limited for other energy sources. However, there is no direct proof that atmospheric energy sources are sufficient to generate *pmf* or drive ATP synthesis. Here, we show that the membrane-associated hydrogen:quinone oxidoreductase Huc from *Mycobacterium smegmatis* enables ATP synthesis from air. Purified Huc couples H_2_ oxidation to the reduction of various ubiquinone and menaquinone analogues. We designed and optimised a minimal respiratory chain in which Huc is reconstituted into liposomes with a *pmf*-generating terminal oxidase and the ATP-generating F_1_F_o_-ATP synthase. Our experiments show that passive hydrogen exchange from air to solution is sufficient for the electron transfer and *pmf generation* required to accumulate ATP. Finally, by combining continuous culture bioenergetics measurements with theoretical calculations, we show this process is sufficient for mycobacteria to sustain *pmf* and ATP synthesis (two ATP molecules per H_2_ oxidised) for maintenance energy requirements during nutrient starvation. These findings prove that atmospheric energy sources are dependable ‘lifeline’ substrates that enable continuous energy conservation during nutrient starvation. In addition, this work provides a new tool for ATP production in synthetic applications, which unlike other approaches is traceless without by-product accumulation.

## Introduction

Metabolic versatility enables bacteria to thrive in diverse ecosystems. Bacteria conserve energy by using a wide range of organic and inorganic energy sources, as well as harvesting light, available in their environments. To do so, they use membrane proteins to transduce chemical and solar energy into a transmembrane electrochemical proton gradient (proton-motive force, *pmf*), driving ATP synthesis through the universally conserved F_1_F_o_-ATP synthase(1, 2). More recently, it was discovered that aerobic bacteria can harness atmospheric energy sources through a process known as ‘aerotrophy’(3, 4). Diverse bacteria produce high-affinity enzymes to consume atmospheric hydrogen (H_2_), carbon monoxide (CO), and sometimes methane (CH_4_)(3, 5–7). Physiological and genetic studies show that this process primarily allows microbes to survive starvation for growth substrates(8–12). This process is globally important, supporting the biodiversity and productivity of soil, marine, and subterranean environments, while regulating the composition of the atmosphere(4, 13–15). Indeed, soil bacteria consume approximately 70 million tonnes of H_2_ from the atmosphere each year, serving as the primary sink in the climate-relevant global hydrogen cycle(16, 17). Yet it remains unclear whether and how this process couples to *pmf* generation and ATP synthesis.

Our recent work has investigated how high-affinity [NiFe]-hydrogenases catalyse atmospheric H_2_ oxidation. The soil bacterium *Mycobacterium smegmatis* uses the group 2a [NiFe]-hydrogenase Huc to selectively oxidize atmospheric H_2_ and is proposed to use the derived electrons for aerobic respiration, allowing it to sustain maintenance needs during organic carbon starvation(9). This impressive feat requires the enzyme to operate at nanomolar affinities (atmospheric H_2_ mixing ratio = 0.53 parts per million)(16), while conserving energy through quinone reduction and preventing inhibition from the high ambient levels of oxygen. The recently solved atomic structure of Huc revealed it forms an octameric complex, with HucSL heterodimers arranged around a tetrameric stalk formed by HucM; this configuration creates a hydrophobic vestibule that allows long-range quinone transport from the bacterial membrane to the quinone-reducing site(6). While this study shows how Huc mediates atmospheric H_2_ oxidation and binds menaquinone, it is unclear how bacteria conserve energy from this process. It has not been demonstrated whether this process is directly coupled to aerobic respiration, generates a *pmf*, or facilitates ATP synthesis. In this study, we addressed these knowledge gaps through combining theoretical, *in vitro*, and *in vivo* studies of the bioenergetics of aerotrophy. By developing a minimal respiratory chain *in vitro*, containing the [NiFe]-hydrogenase Huc, cytochrome *bd-*I oxidase, and ATP synthase, we show that H_2_ oxidation by Huc efficiently drives ATP synthesis, at both elevated and atmospheric H_2_ levels. These findings provide an ultimate biochemical proof that atmospheric trace gases support microbial bioenergetics and enable long-term persistence during nutrient deprivation.

## Results and Discussion

### Huc mediates the exergonic reduction of diverse quinones with atmospheric H_2_

The delivery of electrons from the oxidation of H_2_ to the respiratory chain requires the reduction of respiratory quinones. The [NiFe]-hydrogenase Huc directly reduces menaquinone bound at its small subunit with electrons generated by the oxidation of H_2_ at its active site on the large subunit(6). We used the Nernst equation to calculate how the free energy yield from H_2_ oxidation (E°′_H2/H_^+^ = −414 mV) coupled to menaquinone reduction (E°′_MK/MKH2_ ≈ −74 mV) varies in response to dissolved H_2_ concentration (Supplementary Note 1). Under standard conditions, this process is strongly exergonic, yielding a ΔG°’ of −66 kJ mol^−1^. Even under ambient conditions, where the dissolved H_2_ concentration is 0.42 nM, the ΔG°’ is −30 kJ/mol and thus remains highly energetically favorable (Table S1). Ubiquinones, the other major group of respiratory quinones, have a redox potential that is ∼170 mV higher than menaquinone, rendering their reduction by hydrogenase even more exergonic even though they are absent from mycobacteria. This means that a minimal respiratory chain powered by atmospheric H_2_ will be limited by Huc’s affinity and turnover rate for H_2_ and menaquinone, rather than the thermodynamic driving force for H_2_ oxidation coupled to quinone reduction (Table S1).

On this basis, we tested the capacity of Huc to reduce a set of ubiquinones and menaquinones, which differ in their amphiphilicity due to their different hydrophobic tail length and headgroup structures and redox potentials (Figure 1A). To do so, we spectroscopically followed quinone reduction (20 μM) under anaerobic conditions (to suppress quinone autooxidation) using an enzyme concentration of 5 nM Huc and an excess of substrate (> 500 μΜ H_2_, solution was bubbled with H_2_ for 5 min) (Figure S1). The hydrophilic quinones 1,4-napthoquinone and ubiquinone Q_0_ were poor substrates for Huc compared to the more hydrophobic quinones, with ubiquinone Q_2_ and decylubiquinone (DUQ) being the most active compounds followed by ubiquinone Q_1_ and dimethyl menaquinone (DMNQ) (Figure 1B). Within the set of quinones we tested, activity strongly correlated with increasing quinone hydrophobicity, demonstrated by a plot of activity against calculated logP values (Figure 1B). Depending on the protein preparation, the maximal rate for quinone reduction approached 20 to 40 s^−1^, which is considerably higher than the previously observed maximal rate of 7 s^−1^ determined using the hydrophilic quinone analogue menadione (6). As the most effective co-substrate, we determined the K_m_ value of ∼9 μM for DUQ reduction under Michaelis-Menten conditions (Figure 1C), which is in good agreement with what has been found for other peripheral membrane proteins that interact with the membrane quinone pool, e.g. NDH-2 from *Mycobacterium tuberculosis* (5 μM, (18)) or *Escherichia coli* (18 μM), (19). These measurements provide the first direct observation of quinone reduction by Huc and support the model for how this enzyme drives cellular bioenergetics.

**Figure 1:**
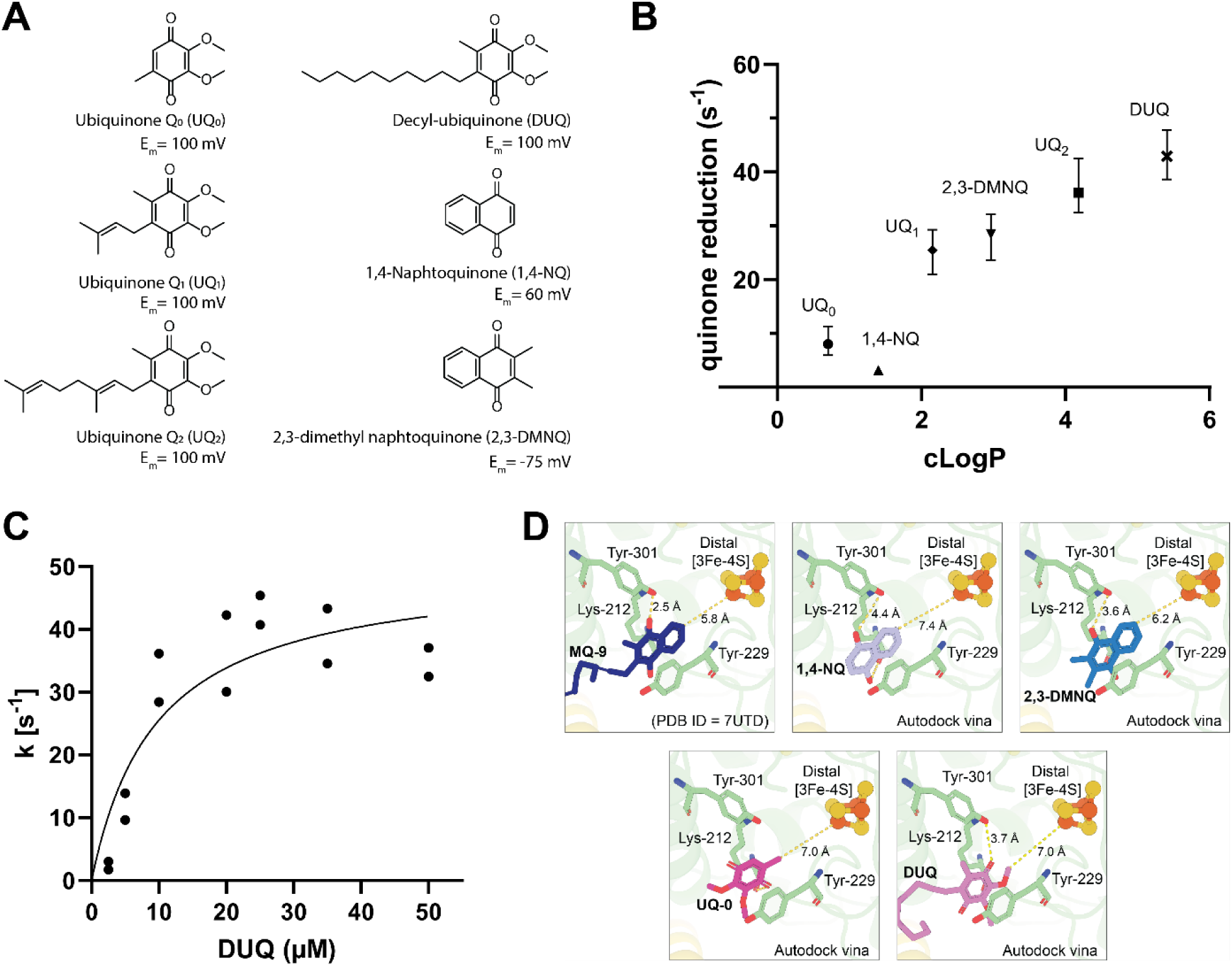
Huc rapidly reduces a various quinones. **A.)** Different ubi- and menaquinones used in this study and their approximate redox potentials are indicated. Potentials were taken from (20–22) **B.)** Rates of quinone reduction of different quinone analogues under Michaelis-Menten conditions plotted against the calculated logP shows a strong activity-hydrophobicity correlation. **C.)** Titration of decylubiquinone under Michaelis Menten conditions. **D.)** Docking simulation of the most relevant quinone analogues with Huc (see text for details).

The more rapid reduction of hydrophobic quinones likely reflects the unique structure of Huc, which contains a long hydrophobic tube that provides a conduit for quinones from the cell membrane to Huc’s internal chamber, containing the electron acceptor site of the small subunits that accommodate and reduce the quinone headgroup. Longer chain hydrophobic quinones such as DUQ are more compatible with Huc’s internal chamber and thus can readily access the quinone-binding site. However, it was surprising that ubiquinones performed comparably to menaquinones in this assay, given that menaquinone is the respiratory quinone in the Huc-producing organism *Mycobacterium smegmatis*, and the structure of Huc was determined with menaquinone bound at its electron acceptor site(6). To investigate this further, we performed *in silico* docking between Huc and the quinones used in the initial activity assays. This analysis indicates that, while the Huc electron acceptor site is better suited to binding the menaquinone (naphthoquinone) head group, it can also accommodate the ubiquinone (benzoquinone) head group (Figure 1D). The energetically more favourable nature of ubiquinone reduction due to its higher redox potential may compensate for their lower structural compatibility in our assay. Finally, it is noteworthy that 1,4-naphthoquinone is a poorer substrate despite having a more energetically favourable midpoint potential (∼100 mV, logP 1.7) compared to DMNQ (−80 mV, logP 3.0), suggesting that substrate hydrophobicity is a more stringent requirement while the active site exhibits plasticity towards different substrates. This is in good agreement with the docking studies showing tighter embedment of the naphthoquinone moiety into the enzyme for DMNQ than 1,4-naphtoquinone, resulting in shorter distances for electron and proton transfer to the [3Fe4S] cluster and Tyr301, respectively (Figure 1D). The finding that Huc can rapidly reduce a both quinone types successfully suggests that the enzyme would be functional in a wide range of respiratory chains, including those dependent on ubiquinone. Moreover, despite not being the physiological quinone for Huc, the experiments suggest DUQ as an ideal electron carrier to reconstitute a Huc-driven minimal respiratory chain powered by atmospheric H_2_.

### Reconstitution of a minimal Huc-powered respiratory chain in liposomes

To couple quinone reduction by Huc to the creation of a *pmf* for ATP generation, we mimicked energy converting membranes by embedding the necessary proteins (quinol oxidase and F_1_F_o_-ATP synthase) into small unilamellar liposomes (Figure 2A). A suitable liposome composition for this system was determined by comparing quinone reduction activity of Huc in the presence of different lipid compositions. As detailed in Supplementary Information 2 and depicted in Figure 2B, Huc loses quickly activity in in the absence of either BSA or liposomes, but did otherwise not vary in activity in response to varying lipid composition, BSA levels, or detergent addition. For all subsequent experiments, we used a liposome composition of PC:PE:PG = 2:2:1, mimicking the overall negative membrane surface of bacteria such as Mycobacteria or *E. coli*.

**Figure 2:**
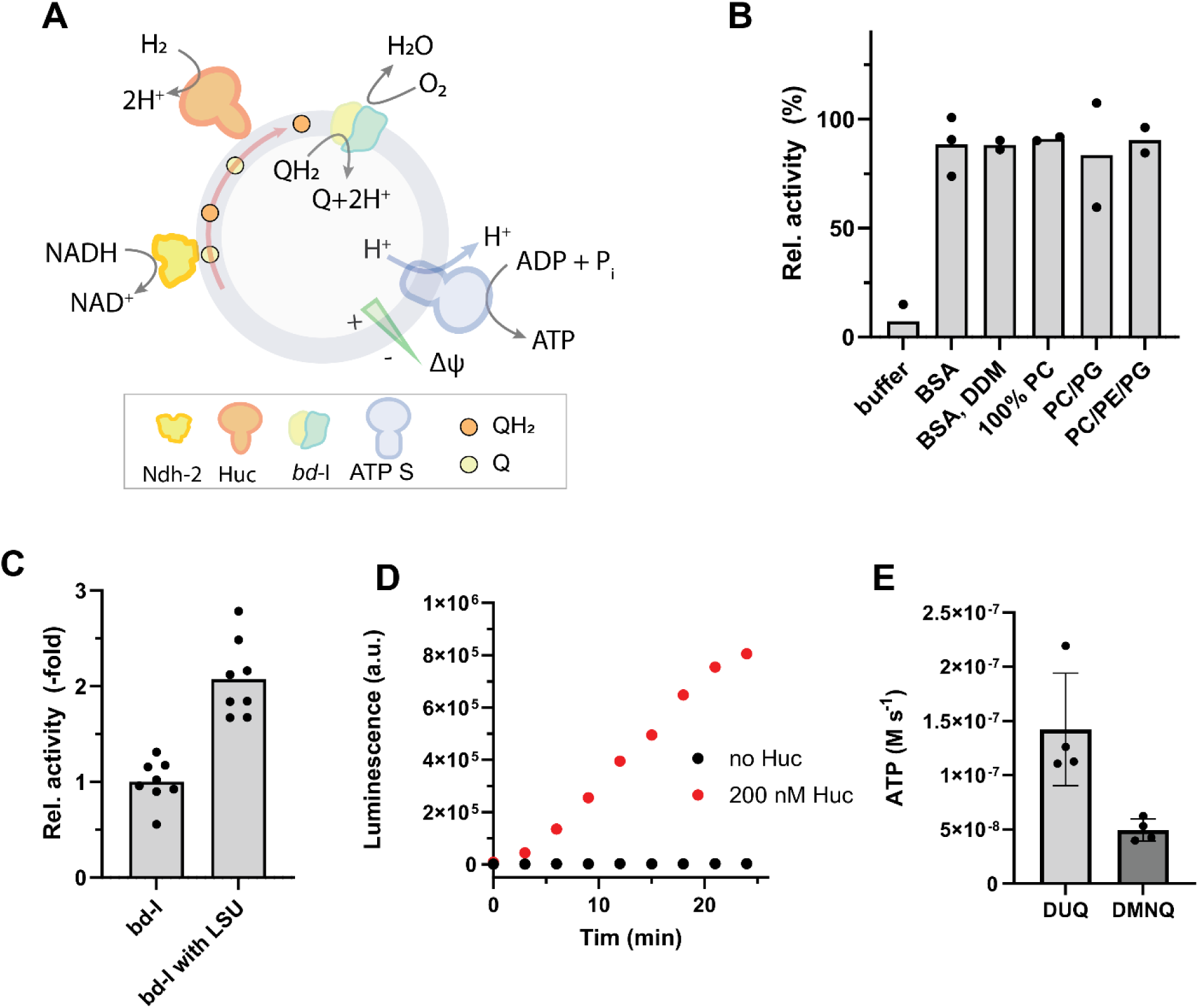
**A.)** Nature-inspired proteoliposomes used in this study. Liposomes containing *bd-*I oxidase and ATP synthase from *E. coli* are energized either using either NADH/NDH-2 or H_2_/Huc using short chain quinones that partition into the liposomal membrane. **B.)** Stability of Huc mediated decylubiquinone reduction under Michaelis-Menten conditions. Different supplementations of the base buffer are indicated. **C.)** Effect of guided *bd*-I insertion during reconstitution into liposomes on ATP synthesis (see Supplementary Note 2 and text for details) **D.)** Raw data from Huc driven ATP synthesis at 100 μM H_2_. Shown are luminescence traces in the absence and presence of 200 nM Huc. **E.)** Summary of four individual reconstitution experiments (as described in D) using either decylubiquinone (DUQ) or dimethylmenaquinone (DMNQ) as electron mediator.

Next, we generated a minimal respiratory chain by reconstituting recombinantly expressed and purified *E. coli* cytochrome *bd*-I terminal oxidase and F_1_F_o_-ATP synthase into these liposomes. We elected to use the *bd* oxidase given it flexibly uses ubiquinone and menaquinone, is present in *M. smegmatis*, and functions well under low-energy settings given its *pmf* formation mechanism; in contrast to canonical heme copper oxidases such as *bo*_3_ or *aa*_3_ oxidase, *bd* oxidases do not pump protons but generate a *pmf* by charge separation (i.e. through the uptake of electrons and protons from opposite sides of the membrane, Figure S2A) (23–25), which conserves less energy but functions also under thermodynamically less favourable conditions, e.g. low oxygen or low quinone/quinol ratio (26, 27). We ensured a functional interaction between the different components showing the detergent solubilized *bd*-I oxidase is capable of rapidly oxidizing the decylubiquinone pool previously reduced by Huc (Figure S2B). Whereas ATP synthase preferably orients uniformly into liposomes (28), the lack of extramembranous components in cytochrome *bd*-I oxidase means it orients more randomly. We therefore transiently modified purified *bd*-I oxidase with a 60 kDa large soluble protein (LSU) (29), promoting a more uniform orientation, leading to an increased *pmf* generation and ATP synthesis rate (Figure S3A). Immediate ATP synthesis (luciferin/luciferase assay) was observed in these liposomes, using dithiothreitol (DTT) and UQ_1_ as a reductant and electron mediator(30, 31), respectively, and rates were two-fold faster in the tagged oxidase variant (Figure 2C and Figure S3B). This system is the first co-reconstitution of a cytochrome *bd*-I oxidase and an ATP synthase in liposome, and these results confirm that electrogenic, but non-pumping *bd* oxidases are sufficient to support cellular ATP synthesis. No drop in ATP synthesis rates was observed within five minutes of operation indicating that a sufficient number of protons is released to the inside of the liposomes by the quinone loop to support continuous proton extrusion by the *E. coli* ATP synthase (10 H^+^ per 3 ATP).

To complete construction of this minimal reconstituted respiratory chain, we replaced the synthetic electron source DTT with electrons from H_2_ liberated by Huc. To achieve this, liposomes containing cytochrome *bd*-I and ATP synthase were preincubated with 50 μM decylubiquinone, ADP, phosphate, and ∼100 μM H_2_ in the headspace of a closed vessel filled with air, and the reaction was started by addition of Huc. Alternatively, Huc was present from the beginning and the reaction was started by addition of ∼100 μM H_2_ via a gastight syringe into the headspace of the vessel. At indicated timepoints, samples were removed, mixed with luciferin/luciferase solution and the ATP levels determined using a luciferin/luciferase assay that reports the dissolved ATP concentration. Immediate and sustained ATP formation was observed during the measurement period, while no ATP is formed in the absence of the Huc enzyme under otherwise identical conditions, indicating Huc is directly driving ATP generation (Figure 2D). Both DUQ and DMNQ could be used as quinone analogues, though rates were two-fold higher with DUQ; this reflects that, in addition to Huc having a higher V_max_ for DUQ reduction (Figure 2E), DMNQ autooxidises ∼5-fold more rapidly under aerobic conditions contributes to this observation (Figure S4A). In parallel experiments, where the peripheral membrane protein NDH-2 of *E. coli* was used to energize the same process, ∼200 times higher rates were found at the same enzyme concentration of 200 nM (Figure S4B), indicating that the Huc-catalysed quinone reduction is the rate-limiting step. Based on this observation, we repeated the above experiment and compared ATP synthesis rates with different amounts of Huc, finding a strictly linear dependency between Huc and ATP synthesis, indicating that concentrations up to 200 nM are not saturating (Figure S4C). Together, these results provide the first direct proof that Huc can couple to terminal oxidases and generate ATP.

### Huc enables ATP synthesis from air in reconstituted liposomes and mycobacterial cells

Given the physiological role of Huc in oxidising atmospheric H_2_, the ultimate test for our minimal electron transport chain was to test if it can produce detectable levels of ATP with atmospheric H_2_ as the sole fuel source. We monitored ATP production by our minimal respiratory chain, using the same experimental setup as previous experiments, but with an open vessel without any additional hydrogen source. As depicted in Figure 3A, ATP synthesis was reproducibly observed. As expected, given the low H_2_ concentrations in the atmosphere (0.53 ppm; corresponding to 0.42 nM in solution), rates were approximately ∼100-fold lower than previous experiments where the dissolved H_2_ concentration was 110 μM. Importantly, no ATP synthesis is observed in the Huc-free control, indicating that H_2_ oxidation by the enzyme is driving ATP synthesis. Moreover, the addition of the potent proton uncoupler SF6847 (Figure 3A) or the ionophores nigericin / valinomycin (Figure S5) strongly affected the rate of ATP synthesis, as expected in a *pmf*-driven system. Interestingly, no discernible difference was seen if either DUQ or DMNQ were used as an electron mediator (Figure 3B). Taken together, the data demonstrate that Huc enables *pmf* generation from air, which in turn drives ATP synthesis.

**Figure 3:**
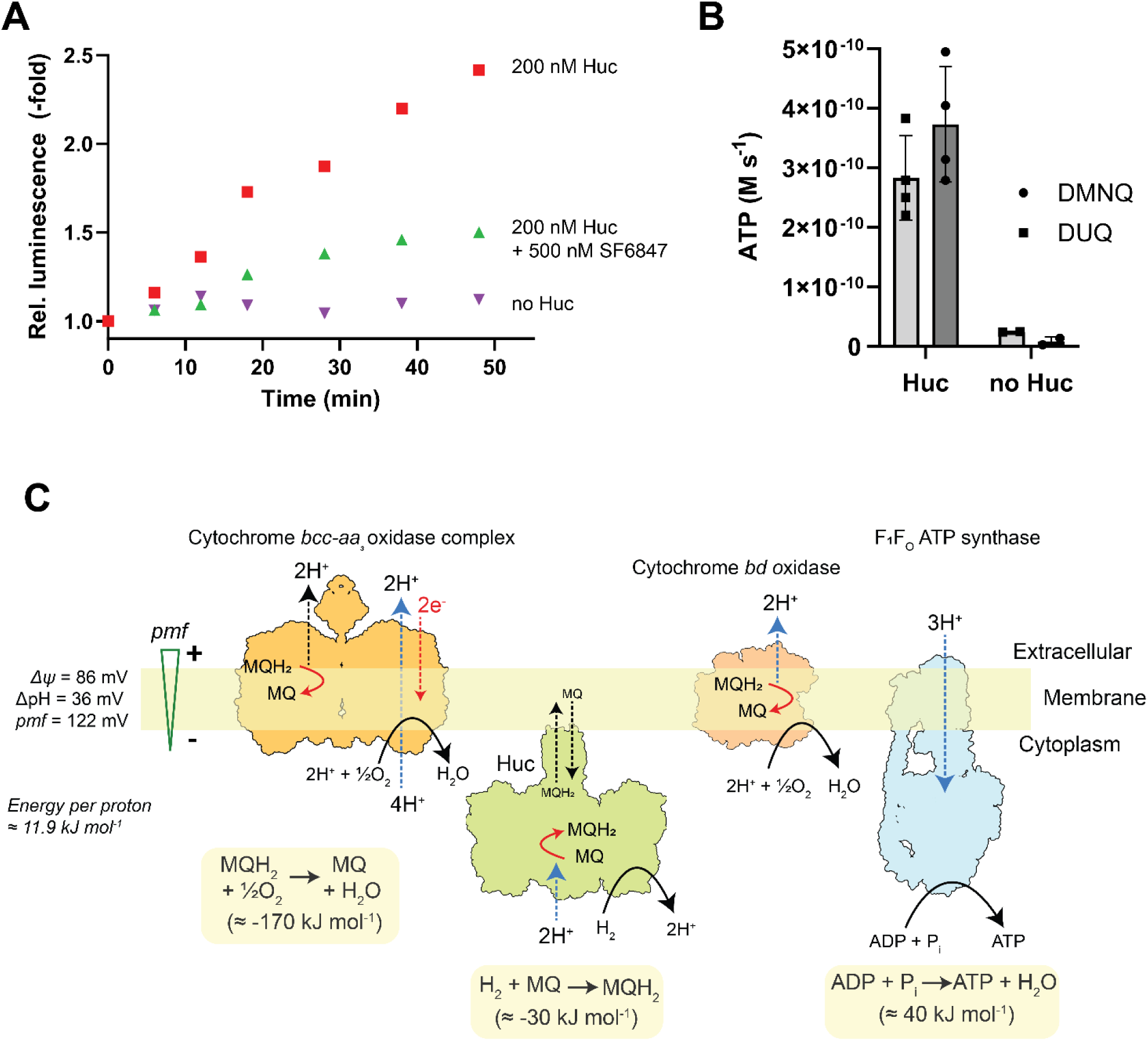
**A.)** Huc driven ATP synthesis from air. Shown are raw traces from three parallel ATP synthesis measurements in an open vessel using 50 μM decylubiquinone as electron mediator, containing either 200 nM Huc (red), 200 nM Huc and 500 nM uncoupler SF6847 (green) or no Huc enzyme (magenta). **B.)** Collected and uncorrected data from biological replicates (n=3) using either DUQ or 2,3-DMNQ as electron mediator in the presence or absence of 200 nM Huc. **C.)** Cartoon display of the bioenergetic scenario for Huc using either *bd-*I oxidase or *bcc*-*aa*_3_ oxidase as *pmf*-generating unit, driving ATP synthesis by the F_1_F_o_-ATP synthase. See text for details.

Having proven that atmospheric H_2_ can be used to produce ATP in a reconstituted system, we finally considered the bioenergetics of Huc-driven ATP synthesis in mycobacterial cells. Based on the Nernst equation (Supplementary Note 1), atmospheric H_2_ oxidation (0.53 ppm) is expected to be highly exergonic when used for aerobic respiration (ΔG°’ = −214 kJ mol^−1^), but also when coupled to the reduction of the other mycobacterial terminal electron acceptors nitrate (ΔG°’ = −125 kJ mol^−1^) and fumarate (ΔG°’ = −50 kJ mol^−1^), using the standard potential at pH 7 for the electron acceptors (Table S1). As oxidation of hydrogen and reduction of menaquinone by Huc is non-electrogenic and no energy is conserved in form of a *pmf*, the available ΔG for ATP synthesis is reduced by ∼30 kJ mol^−1^. Next, we considered the number of protons translocated per molecule of H_2_ oxidized (i.e. H^+^ / 2e^−^ ratios) when electrons are transferred to the two terminal oxidases of mycobacteria, the cytochrome *bd* oxidase and cytochrome *bcc*-*aa*_3_ oxidase super complex. For the *bd* oxidase, given cytosolic proton consumption by terminal oxidases (2H^+^/2e^−^) is counterbalanced by conversion of H_2_ to protons by Huc, proton translocation is expected to occur exclusively through a scalar mechanism *via* the menaquinone redox loop (i.e. 2 charges translocated and 2H^+^ released upon quinol oxidation by *bd* oxidase per H_2_ molecule oxidized, see Figure S2A and 3C). In contrast, proton pumping by the cytochrome *bcc*-*aa*_3_ oxidase (32) would greatly increase efficiency (i.e. 6 H^+^ translocated per H_2_ molecule oxidized, Figure 3C), with our physiological experiments suggesting Huc efficiently couples to this oxidase despite potential backpressure(33). As mycobacteria encode an efficient F_1_F_o_-ATP synthase with a *c*-ring stoichiometry of 9(34, 35), they theoretically require three protons per ATP molecule produced, and thus one molecule of atmospheric H_2_ can be used to produce either ^2^/_3_ or 2 ATP depending on the terminal oxidase used when coupling is optimal. The latter value is compatible with the thermodynamics of the reaction of menaquinol with oxygen (ΔG= −170 kJ mol^−1^) and the ΔG value for cellular ATP synthesis (ΔG°’ ∼ 40 kJ mol^−1^). In comparison, mitochondria have similar thermodynamics (NADH + ½ O_2_ + H^+^ è NAD^+^ + H_2_O, −214 kJ/mol), but with an additional four proton pumped by complex I (10 H^+^ per NADH) and a *c*_8_ ring of the ATP synthase yielding ∼3 ATP per NADH oxidized(1).

Given these theoretical considerations, we next profiled the cellular energetics of continuous cultures of *M. smegmatis*, to determine whether atmospheric H_2_ oxidation is sufficient to generate ATP for maintenance energy requirements (e.g. *pmf*) Figure S6 and Table 1). Steady-state cultures were grown on minimal salts media supplemented with limiting concentrations of glycerol, at dilution rates ranging from 0.02 h^−1^ (doubling time 35 h, very slow growth) to 0.14 h^−1^ (doubling time 5 h, fast growth). As detailed in Supplementary Note 3, glycerol consumption rates, cell yield (CFU mL^−1^), and cell yield per mol glycerol utilized and ATP consumed (Y_glycerol_, Y_ATP_) were each linear functions of dilution rate (Figure S6A-D). With respect to ATP synthesis, atmospheric H_2_ oxidation is insufficient to meet the relatively high ATP requirements of this microorganism for growth (i.e. Y_ATP_) (Table 1). These Y_ATP_ values are equivalent to an ATP demand of 263 mmol ATP per gram of newly synthesized biomass, a demand for ATP that is 3 to 4-fold higher than those reported for glycerol- or glucose-limited cultures of other aerobic bacteria (54, 55) suggesting an important requirement for high ATP yielding metabolism (ie. oxidative phosphorylation) in the growth of *M. smegmatis*. In contrast, the the rate of ATP utilization for maintenance functions (mATP) was 3 mmoles of ATP/h/g [dry weight] of cells and therefore atmospheric H_2_ oxidation would appear sufficient to provide ATP for maintenance functions given the high predicted ATP yield per H_2_ molecule oxidised (Figure S3C). We calculated the *pmf* of these cultures (by measuring and summing the membrane potential and pH gradient components of *pmf*), yielding values between 122 to 116 mV (Table 1). On this basis, we calculate ∼12 kJ mol^−1^ is required to transport each proton across the membrane (1), accumulating ∼36 kJ/mol per 3 H^+^ and ATP synthesized, being well in accordance with a ΔG ∼34 kJ/mol for ATP synthesis at a ATP/ADP ratio of ∼1 (1). Under fast-growing conditions, the ATP/ADP ratio is typically higher (>10), yielding a higher ΔG and thus requiring a larger Δψ if only 3 H^+^ per ATP are pumped(36). With an energy of −170 kJ/mol at hand (based on the redox couple of menaquinol to O_2_), enough thermodynamic driving force is present to translocate 6 H^+^ as proposed for the *bcc*_1_-*aa*_3_ supercomplex required to synthesize 2 ATP as discussed above. The ability to also use a *bd* complex enables the cell to maintain ATP production from air, albeit with lower efficiency, even in response to varying energy conditions. Altogether, this demonstrates that *M. smegmatis* can continually maintain *pmf* and generate ATP through reacting H_2_ and O_2_ from the atmosphere using minimalistic respiratory chains, with similar configurations to the one reconstituted in this study. Our calculations suggest that ATP synthesis is thermodynamically feasible and that binding and oxidation of H_2_ by Huc is the rate limiting step, agreeing with our previous work showing strong upregulation and activity of high-affinity hydrogenases, together with the severe survival phenotypes when knocked out, under energy-limited (including glycerol-starved) conditions(9, 10, 37).

**Table 1:**
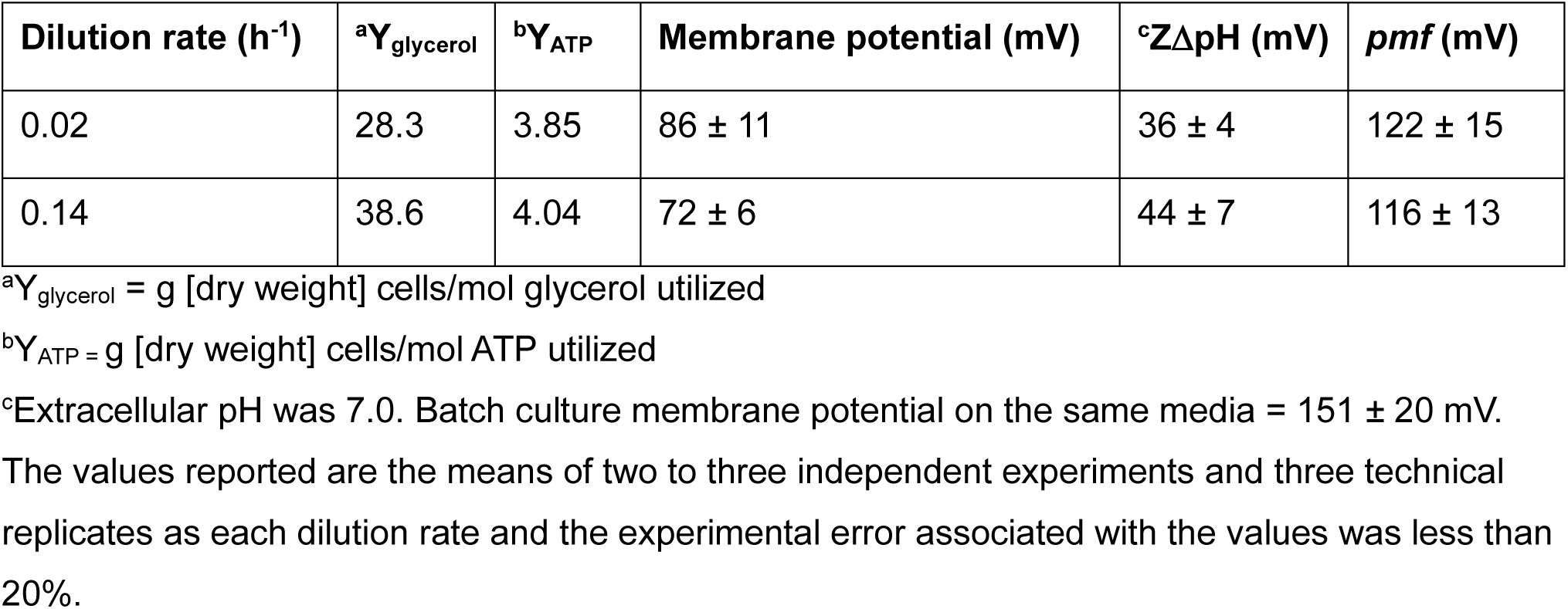
Bioenergetic parameters measured at steady state in glycerol-limited continuous culture of *M. smegmatis*.

## Conclusions

This work proves that aerotrophy is a viable strategy for microbial life. We provide critical theoretical and experimental support for the hypothesis that atmospheric energy sources are sufficient to sustain microbial survival. By reconstituting a minimal nature-inspired respiratory chain, we show that Huc-mediated atmospheric H_2_ oxidation can drive quinone reduction and generate ATP. Moreover, by combining theoretical calculations with bioenergetic measurements, we show that the coupling of these processes in mycobacterial cells is sufficient to sustain their *pmf* and generate 2 ATP per H_2_ molecule under highly energy-limiting conditions. Such findings help to rationalise previous physiological studies showing high-affinity hydrogenases are among the most expressed, active, and important (based on phenotypes when deleted) enzymes for mycobacteria and other species during energy-limiting conditions(8–12, 37–40). Indeed, while the concentrations of atmospheric H_2_ are modest, this omnipresent substrate is still sufficient to drive much *pmf* production and ATP synthesis, thereby providing a dependable lifeline for long-term survival. It should be noted that, while mycobacteria appear to primarily use high-affinity hydrogenases for survival, these enzymes also enhance growth on highly oxidised organic substrates, at limited substrate concentrations (as per the continuous culture measurements), or when H_2_ is available at elevated levels, with the considerable *pmf* and ATP provided by H_2_ oxidation providing a modest but considerable advantage(10, 37, 41).

More broadly, these findings help to explain why high-affinity hydrogenases are so widely distributed and active in nature(14, 17). Numerous environmental microbes will benefit from the reliable supply of H_2_ at atmospheric concentrations and often elevated levels through biological, geological, and anthropogenic processes(3). While the ATP demands of *M. smegmatis* are too steep for atmospheric H_2_ oxidation to meet catabolic requirements during growth, some microbes that either have low energy demands (e.g. the ultramicrobacterium *Sphingopyxis alaskensis*(15)) or rely on high-potential substrates (e.g. nitrite, sulfide, or iron oxidisers(39)) meet their energy needs by constitutively consuming this gas alongside other substrates, with a few microbes even appearing to grow on air alone (e.g. *Methylocapsa gorgona*)(7, 13, 42). Altogether, the numerous microbes that use atmospheric H_2_ for growth and survival drive a large, robust sink in the global hydrogen cycle, consuming 70 teragrams of this gas per year(43). Finally, we note that these findings have potential biotechnological applications. High-affinity oxygen-insensitive hydrogenases, in combination with ATP synthase modules, may be applied to create energy-autonomous cell-based and cell-free synthetic biology platforms(44–46). Gutierrez et al. demonstrated ATP synthesis using the hydrogenase from *Desulfovibrio vulgaris* under anaerobic conditions with 1 atm hydrogen, using a gold electrode as an electron acceptor; proton release acidified the compartment, establishing a pH gradient to drive ATP synthesis in a planar bilayer setup with reconstituted ATP synthase (33, 34). The liposome-based system presented here, requiring only ambient substrates, could be more readily integrated into synthetic biology applications especially where anoxic conditions, elevated substrates, and external electron acceptors may be impractical. A further advantage of this process, for microbes in nature and for biotechnology alike, is that the only end-product of this reaction is water.

## Material and Methods

### Chemicals

If not otherwise stated, chemicals were obtained from Sigma/Merck. Ubiquinols Q_1_, Q_2_, DUQ and menaquinol DMNQ were synthesized as described(47).

### Enzyme expression and purification

#### Huc

*Mycobacterium smegmatis* mc^2^155 PRC1 HucS-2×StrII cells were cultured at 37 °C, 150 rpm, for five days into the stationary phase in 24 L of Hartmann’s de Bont (HdB) minimal medium (per litre: 2 g (NH_4_)_2_SO_4_, 1 g NaH_2_PO_4_**·**H_2_O, 1.55 g K_2_HPO_4_, 10 mg Na_2_EDTA**·**2H_2_O, 1 g MgCl_2_**·**6H_2_O, 1 mg CaCl_2_**·**2H_2_O, 0.2 mg Na_2_MoO_4_**·**2H_2_O, 0.4 mg CoCl_2_**·**6H_2_O, 1 mg MnCl_2_**·**4H_2_O, 2 mg ZnSO_4_**·**7H_2_O, 5 mg FeSO_4_**·**7H_2_O, 0.2 mg CuSO_4_**·**5H_2_O, 1 mg NiSO_4_**·**6H_2_O, 2.5 mL Tyloxapol, 1 mL Glycerol). Cells were harvested by centrifugation and resuspended in lysis buffer (50 mM Tris, 150 mM NaCl, pH 8.0) supplemented with 0.1 mg mL^−1^ lysozyme, 0.1 mg mL^−1^ DNase, 1 mM MgCl_2_, and protease inhibitor cocktail tablets (Merck). Lysis was performed using a cell disruptor (Emulsiflex C-5), and cell debris was removed twice by centrifugation at 30,000 g for 30 min. Clarified lysates were treated with biotin blocking solution (1 mL per 50 mL lysate; IBA Lifesciences) before being loaded onto a 1 mL StrepTrap HP column (Cytiva). The column was washed extensively with lysis buffer and bound Huc was eluted using lysis buffer containing 2.5 mM desthiobiotin. Huc-containing fractions were identified by SDS-PAGE, pooled, and concentrated using a 100 kDa MWCO centrifugal concentrator (Amicon, Millipore). The concentrated Huc was further purified by size exclusion chromatography using a Superose 6 10/300 GL column (Cytiva). Fractions containing oligomeric Huc (i.e. HucLSM subunits were observable on SDS-PAGE) were pooled, concentrated to >5 mg mL⁻¹, flash-frozen in liquid nitrogen, and stored at −80 °C. The typical yield of purified Huc per litre of culture was 100-300 μg L⁻¹.

#### NDH-2, F_O_F_1_ ATP synthase and MBP-SpyCatcher

NDH-2, F_O_F_1_ ATP synthase and MBP-SpyCatcher were expressed and purified as described previously(*29, 31*). NDH-2 and F_O_F_1_ ATP synthase were stored in Ni-NTA elution buffer (10 mM HEPES pH 7.4, 100 mM NaCl, and 10 mM KCl, 200 mM imidazole, 0.05% DDM for NDH-2), (50 mM MOPS-NaOH pH 8, 100 mM NaCl, 5 mM MgCl_2_, 30 g/l sucrose, 10 % glycerol, 285 mM imidazole, 0.005 % LMNG for F_O_F_1_ ATP synthase). After affinity chromatography, MBP-SpyCatcher was further purified with gel filtration in PBS. All proteins were divided into aliquots, flash frozen in liquid nitrogen and stored at −80°C before use.

#### Cytochrome *bd-I oxidase*

*bd*-I oxidase was purified using plasmid pET28b(+)*cydA*_his_*BX* (48), coding for subunits A, B and X with a hexahistidine tag at the C-terminus of the A subunit, with immobilized metal affinity chromatography as described by Thessling *et al*(23). The Ni-NTA column was eluted with 50 mM MOPS pH 7, 500 mM NaCl, 0.003% LMNG (Anatrace, Maumee, US) using an imidazole gradient from 20 – 500 mM over 3 cv. The peak fractions were pooled, concentrated (Amicon Ultra, 100 kDa cut-off, Merck, Darmstadt, Germany), aliquoted, flash frozen in liquid nitrogen and stored at −80°C.

### Quinone reduction

Enzymatic quinol reduction measurements were performed in buffer containing 50 mM tris-HCl pH 8, 150 mM NaCl, 0.3 mg mL^−1^ Bovine Serum Albumin that was flushed with nitrogen (Carbagas, Gümligen, Switzerland) for 20 minutes and saturated with hydrogen (Carbagas, Gümligen, Switzerland) for further 10 minutes in a sealed quartz cuvette (Hellma Analytics, Müllheim, Germany). If not otherwise indicated, 50 µM of the respective quinone (from a 10 mM stock solution in dry EtOH) was added to the buffer above and equilibrated for 1 minute, before the reaction was started by addition of 5 nM Huc. Subsequent quinone reduction was monitored with a Cary 60 UV-Vis spectrometer (Agilent Technologies, Santa Clara, US). The quinones were measured at the specific absorption maxima: Ubiquinone-0 (UQ_0_) at 268 nm, Ubiquinone-1 (UQ_1_) at 275 nm, Ubiquinone-2 (UQ_2_) 275 nm, Decylubiquinone (DUQ) at 278 nm, 1,4-Naphthoquinone (1,4-NQ) at 246 nm, and 2,3-Dimethyl-1,4-naphthoquinone (2,3-DMNQ) at 270 nm.

### Quinol autooxidation

DUQ and 2,3-DMNQ stocks were prepared in argon saturated anhydrous EtOH. Crystals of sodium borohydride were added until the solution was completely transparent. After 15 minutes of incubation on ice, 5M HCl was added in 1 µl steps until bubble formation stopped. The white precipitate was removed by centrifugation (10,000 g, 10 min, 4 °C) and clear supernatant was separated, flushed with argon (Carbagas, Gümligen, Switzerland) and stored at −20°C. Reduced DUQ and 2,3-DMNQ were diluted to 10 µM in 50 mM tris-HCl pH 8, 150 mM NaCl and autooxidation at aerobic conditions was measured at the respective absorption maximum of the quinones (288 nm for DUQ and 270 nm for 2,3-DMNQ) in a quartz cuvette (Hellma Analytics, Müllheim, Germany) using a Cary 60 UV-Vis spectrometer (Agilent Technologies, Santa Clara, US). Finally, quinols were completely oxidised by addition of 15 nM *bd*-I oxidase.

### Molecular docking of quinone analogues

To assess the ability of 1,4-NQ, 2,3-DMNQ, UQ-0, and DUQ to bind at the Huc MQ-9 binding electron acceptor site, we used the Autodock Vina via the command line (49, 50). Models for 1,4-NQ, 2,3-DMNQ, UQ-0, DUQ, and Huc (PDB ID = 7UTD) were obtained from the PDB and prepared for docking using Autodock Vina ligand preparation tools. A search box was set encompassing the Huc electron acceptor binding site and a total of nine binding modes were sought for each docking run, with search exhaustiveness of 300 and a maximum energy difference of 3 kcal per mol. The Autodock Vina force field was used for docking (49, 50).

### Detergent-mediated co-reconstitution of *E. coli bd*-I oxidase and F_O_F_1_ ATP synthase

#### Liposome preparation

1,2-dioleoyl-sn-glycero-3-phosphocholine [18:1 (Δ9-cis)] (DOPC), 1,2-dioleoyl-sn-glycero-3-phosphoethanolamine [18:1 (Δ9-cis)] (DOPE) and 1,2-dioleoyl-sn-glycero-3-phospho-(1’-rac-glycerol) [18:1 (Δ9-cis)] (DOPG) were obtained from Avanti Polar Lipids (Alabaster, US). Chloroform stocks of the lipids were mixed in the desired ratios. The mixture was flushed with argon gas (Carbagas, Gümligen, Switzerland) to evaporate the chloroform and spread the lipids in a thin layer. The thin lipid film was dried in a desiccator for at least two hours, but usually overnight. Lipids were then rehydrated in 20 mM HEPES pH 7.4, 2.5 mM MgCl_2_, 50 g/L sucrose and subjected to seven freeze-thaw cycles (29.4 °C / LN_2_). Before reconstitution, the lipid mixture was extruded 17 times through a 200 nm Whatman filter (Cytiva, Marlborough, US). Soybean lecithin containing 90% PC (Alfa Aesar, Ward Hill, US) was obtained as powder and directly solubilized in the desired buffer by vortexing. Once homogeneous the mixture was subjected to seven freeze-thaw cycles (29.4 °C / LN_2_). Before reconstitution, the lipids were extruded at least 17 times through a 100 nm Whatman filter (Cytiva, Marlborough, US).

#### bd-I oxidase orientation via guided insertion during detergent-mediated reconstitution

Before reconstitution, *bd*-I oxidase was coupled to *tris*NTA-MBP acting as a large soluble unit (LSU) as described in Amati *et al*(29). MBP-SpyCatcher protein was incubated with a 3-fold excess of the bifunctional *tris*NTA-SpyTag linker and incubated for 2 hours at 25 °C and 1200 rpm. Excess *tris*NTA-SpyTag was removed by a prepacked gel filtration column (CentriPure P10, emp Biotech, Berlin, Germany). The resulting *tris*NTA-MBP was then coupled to the HisTag displayed by the *bd*-I oxidase before reconstitution by incubation with a 5-fold molar excess of *tris*NTA-MBP (over *bd*-I oxidase) and 0.5 mM NiSO_4_. The mixture was incubated for 5 minutes at 25 °C.

#### Detergent mediated co-reconstitution into preformed liposomes

Preformed liposomes (10 mg/ml) were destabilized with 0.6 % Na-cholate. Approximately 5 F_1_F_o_-ATP synthases and 10 *bd*-I oxidases per vesicle were added and the mixture was incubated for 30 minutes at room temperature with occasional gentle flicking. Detergent was removed by gel filtration using a CentriPure 10 – Z25M column (emp Biotech, Berlin, Germany) preequilibrated with 20mM HEPES pH 7.4, 2.5 mM MgCl_2_, 50 g/L sucrose.

### ATP synthesis

#### Driven by Q_1_/DTT

Liposomes containing F_1_F_o_-ATP synthase and *bd*-I oxidase were diluted to 0.3 mg/ml in ATP synthase measurement buffer (20 mM Tris-PO_4_ pH 7.4, 5 mM MgCl_2_), containing 4 mM DTT, 0.2 mM ADP and 0.4 mg/ml luciferase/luciferin solution (ATP Bioluminescence Kit CLS II; Roche, Basel, Switzerland). The luminescence signal was measured with a GloMax® 20/20 Luminometer (Promega, Madison, US) and the measurements were normalised by addition of 0.1 µM ATP in the beginning of the measurement. The reaction was started by addition of 15 µM ubiquinone Q_1_.

#### Driven by NDH-2/NADH

Liposomes containing F_O_F_1_ ATP synthases and *bd*-I oxidase were diluted to 0.6 mg/ml in ATP synthase measurement buffer (20 mM Tris-PO_4_ pH 7.4, 5 mM MgCl_2_), containing 0.2 mM ADP, 50 µM DUQ, 1 mM NADH (Santa Cruz Biotechnology, Dallas, USA) and 0.4 mg/ml luciferase/luciferin solution. The luminescence signal was measured with a GloMax® 20/20 Luminometer (Promega, Madison, US) and the measurements were normalised by addition of 0.1 µM ATP at the beginning of the measurement. The reaction was started by addition of 200 nM NDH-2.

#### Driven by Huc/H_2_

Liposomes containing F_1_F_o_-ATP synthase and *bd*-I oxidase were diluted to 0.6 mg/ml in 20 mM Tris-PO_4_ pH 7.4, 5 mM MgCl_2_, 0.2 mM ADP, 50 µM DUQ or DMNQ. The mixture was prepared in a sealable 4.5 ml cuvette (Hellma Analytics, Müllheim, Germany) and constantly stirred. To start the reaction 700 µl hydrogen (Carbagas, Gümligen, Switzerland) and 200 nM Huc were added with a gastight syringe (Hamilton, Bonaduz, Switzerland). ATP synthesis at atmospheric conditions was performed as described above, but in an open vessel with an air/liquid interface of ∼5 cm^2^. The reaction was mildly stirred. At defined timepoints, 25 µl of the reaction was mixed with 75 µl 500 mM Tris-HCl pH 7.75, containing 0.4 mg/ml luciferine/luciferase (ATP Bioluminescence Kit CLS II; Roche, Basel, Switzerland). The luminescence signal was measured with a GloMax® 20/20 Luminometer (Promega, Madison, US). Every measurement was normalised by addition of 0.4 µM ATP and repeated luminescence measurement.

### Continuous culture

*Mycobacterium smegmatis* strain mc^2^155 was grown with agitation at 37°C in either lysogeny broth supplemented with 0.05% (w/v) Tween 80 (Sigma chemicals) (LBT) or Middlebrook 7H9 broth (Difco Laboratories, Detroit, Mich.) supplemented with sterile Middlebrook ADC enrichment (Becton Dickinson, Cockeysville, Md.). For solid media, Middlebrook 7H11 was supplemented with ADC and glycerol (0.5% v/v) or LBT with 1.5% agar. Middlebrook 7H9 basal medium supplemented with 0.1% (w/v) Tween 80 and 0.2% (w/v) glycerol was used to culture *M. smegmatis* mc^2^155 continuously in a New Brunswick BioFlo fermenter model C30 using a 350 ml constant culture volume at 37°C. The culture system was controlled by nutrient addition from the medium reservoir by use of a LKB peristaltic pump. The pH of the growth medium remained at pH 7.0 ± 0.2 during continuous culture at the dilution rates used in this study. The growth medium was inoculated with exponential-phase cells and permitted to undergo batch growth (no medium feed rate, and consequently no dilution rate, was applied during this time) to densities of approximately 0.3-0.8 (OD_600_) with constant impeller agitation at 240 rpm. Concomitant with initiating continuous growth, agitation was increased to 400 rpm to provide adequate aeration during steady state conditions. Each dilution rate was maintained at a stable optical density, pH level and aeration for at least four residence times to allow the culture to reach steady state prior to sampling. Chemostat cultures were routinely checked for potential bacterial and fungal contamination by plating samples onto LBT agar during the experiment.

#### Bioenergetic measurements

At each steady state, 100 mL of culture was removed for the following analyses: residual glycerol and end-product analyses, total viable colony forming units (CFU) and cellular dry weight determinations. The values reported are the means of two to three independent experiments that were performed in technical triplicate, and the experimental error associated with the values was less than 20%. Residual glycerol was determined by assay with glycerol dehydrogenase measuring NAD^+^ reduction at 340 nm with a Cary 50 (Varian) spectrophotometer. Cell dry weight was measured by harvesting (14,000 rpm, 10 min, 4°C) and washing cells from 25 mL of steady state culture onto a 0.45 μm membrane filter [Millipore (47 mm)] and drying at 55°C to a constant weight. The filter was cooled in a desiccator for 15 min before being weighed. End-products from bacterial growth were measured by gas chromatography using a Hewlett-Packard HP5890 series II gas chromatograph fitted with a wide-bore carbowax capillary column (Econocap 19659, Alltech Associates) using conditions and procedures as described previously (51). Cell viability was determined by colony growth on LBT agar plates. Protein concentrations were determined using a bicinchoninic acid protein assay kit (Sigma), with bovine serum albumin as the standard. The membrane potential and transmembrane pH gradient (ΔpH) were determined in chemostat grown cells as previously described (52). The internal pH was determined from the distribution of [^14^C]benzoate (10–25 mCi mmol^1^, 11 µM) using the Henderson-Hasselbalch equation. The ZΔpH was calculated as 62 mV × ΔpH. The membrane potential was calculated from the uptake of [^3^H]methyltriphenylphosphonium iodide ([^3^H]TPP^+^, 5 µM) according to the Nernst relationship with corrections for non-specific binding. (52)

## Footnotes

### Author contributions

C.v.B., C.G., and R.G. conceived and designed this project. S.S. and S.U.M. performed quinone and liposome experiments, J.P.L. and A.K. provided Huc preparations. G.M.C. and S.T. performed mycobacterial bioenergetic measurements. R.G. performed docking simulations. C.v.B., C.G., R.G., G.M.C., S.S., and S.U.M. analysed data. C.v.B., C.G., R.G., G.M.C., and S.U.M. wrote the manuscript with input from all authors.

## Acknowledgements

We thank Dr. Alexander Thesseling and Prof. Thorsten Friedrich (University of Frankfurt) for plasmid pET28b(+)*cydA*_his_*BX.* Work in the CvB lab is supported by Swiss National Science Foundation and University of Bern research Foundation. This work was supported by ARC Discovery Grants (DP230103080, DP200103074) awarded to C.G. and R.G., NHMRC Emerging Leadership Fellowships (APP1178715 to C.G.; APP1197376 to R.G.), an ARC Future Fellowship (FT240100502 to C.G.) and a Lottery Health Grant awarded to G.M.C. We thank Philipp Grossenbacher and Martin Lochner (University of Bern) for synthesis of ubiquinones Q_1_, Q_2_, DUQ and 2,3-DMNQ.

## Conflicts of interest

The authors declare no conflicts of interest.

## Supplementary Information

### Supplementary note 1. Calculations of the energy yield from atmospheric H_2_

Thermodynamically, the oxidation of hydrogen

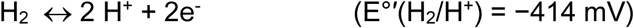

coupled to the reduction of menaquinone

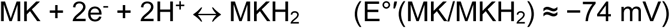

is highly exergonic. Using

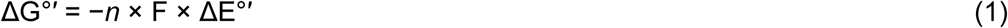

where *n* = 2, equaling the number of electrons transferred, F is the Faraday constant (96.5 kJ/(V x mol)) and ΔE°′ is the difference between the two redox potentials (−414 mV – (−74 mV) = −340 mV), a ΔG°’= −65.5 kJ/mol is obtained.

To participate in chemical reactions with hydrogenases such as Huc, however, hydrogen must be dissolved in the aqueous environment of bacterial cytoplasm or the buffer solution containing the purified enzyme. Hydrogen has low to moderate solubility in water (0.6 fold compared to oxygen), and at room temperature and 1 atm, up to 800 μM can be dissolved according to Henry’s law using a k_H_ = 7.8 × 10^−4^ mol/(L × atm). However, atmospheric hydrogen is only present at 0.53 ppm, corresponding to an aqueous concentration of 0.42 nM. This low concentration does not only necessitate a very affinity of the binding site for hydrogen (threshold ≤ 0.031 nM) as discussed in Grinter et al (6), but also have a direct impact on the thermodynamics of the reaction as the correct redox potential has to corrected for the actual substrate and product concentration. Assuming an overall air pressure of 1 atm, the partial pressure of H_2_ in air is 0.53 × 10^−7^ atm

Applying the Nernst equation (2) to for the H_2_/H^+^ couple yields

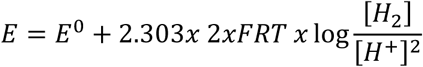

where R is the gas constant and, and at pH 7, where [H^+^] is 100 nM and [H_2_] is 0.53 x 10^−7^ atm, obtaining an adjusted midpoint potential of −229 mV, leading to a an actual ΔG°’ = −29.9 kJ/mol for the reaction with menaquinone, which is still strongly exergonic, reflecting that the reaction is not thermodynamically limited but rather by the requirement of the high hydrogen affinity site. In addition, reduced menaquinone has to be protected from autooxidation in the presence of high oxygen concentrations as experienced for *M. smegmatis*.

Find below a list of further ΔG values that have been calculated using equations (1) and (2) and are mentioned in the text.

**Table S1:** Thermodynamic parameters of electron donors and acceptors.

| Redox couple | $E_{m,7}$ (pH 7, 25° C)<br>(mV) | $\Delta G$ with atmospheric<br>$H_2$ (kJ/mol) | $\Delta G$ for reaction with<br>menaquinol (kJ/mol) |
| --- | --- | --- | --- |
| $H_2/H^+$, standard (1 atm) | -414 | | |
| $H_2/H^+$, atmosph. ($0.5 \times 10^7$ atm) | -229 | | |
| Menaquinone/ol | -75 | -29.9 |  |
| Ubiquinone/ol | +100 | -63.5 |  |
| $O_2/H_2O$ (1 atm $O_2$) | +820 | -202.4 | -172.5 |
| $NO_3^-/NO_2^-$ | +420 | -125.2 | -95.3 |
| Fumarate/succinate | +30 | -50.0 | -20.1 |

### Supplementary Note 2. Optimisation of liposome composition

We tested how varying liposome composition affected Huc activity. We previously observed that the addition of bovine serum albumin (BSA) was important for stabilising Huc at low concentrations, likely due to molecular crowding. Our current analysis is consistent with this finding, with the addition of 0.3 mg mL^−1^ BSA required to stabilise the enzyme, with no further improvement at a BSA concentration to 1.5 mg mL^−1^ or detergent (Figure 2B). Interestingly, in the presence of empty liposomes, supplementation of BSA was not necessary to maintain Huc activity which remained stable during the measurement period. No discernible difference in activity was found in the presence of liposomes containing only zwitterionic PC liposomes, or a mixture of PC or PC/PE and the negatively charged PG liposomes (Figure 2B). The terminus of the Huc tube structure responsible for membrane interaction contains a number of lysine and arginine residues, which give the region a net positive charge(6). As such, a negatively charged liposome surface is expected to be necessary for interaction with Huc, which is consistent and the negatively charged surface of the bacterial membrane (53). To ensure conditions for optimal protein-membrane interaction, a PC:PE:PG = 2:2:1 was used in all future experiments.

### Supplementary Note 3. Bioenergetics of energy-limited *Mycobacterium smegmatis* cultures

We measured the bioenergetic parameters of steady-state glycerol-limited continuous cultures (37) of *M. smegmatis* grown over a dilution range of 0.02 to 0.14 h^−1^, which represents a doubling time of 35 (very slow growing) to 5 h (fast growth) respectively. Under these conditions, the glycerol consumption rate (mmol of glycerol consumed/h/g [dry weight cells]) was a normal linear function of the dilution rate (Figure S6). Under these conditions, glycerol was oxidized to CO_2_ at each dilution rate (i.e. no partial oxidation occurred and no end-products, e.g. acetate or lactate, were detected in the growth medium). The slope of the glycerol consumption rate is equivalent to 1/Y_glycerol_^max^ and the coordinate intercept is the glycerol consumption rate for maintenance (m_glycerol_) energy requirements. Based on these calculations, *M. smegmatis* exhibited a Y_glycerol_^max^ of 40 g [dry weight] cells/mol glycerol utilized and an m_glycerol_ of 0.28 mmol of glycerol/h/ g [dry weight] cells (Figure S6B). The cell counts (CFU mL^−1^) were 1 × 10^7^ to 1 × 10^8^ over the dilution series studied (Figure S6A).

On the basis of the molar growth yield and assuming the carbon content of the cells is 50% (assimilated into cell biomass), we applied the following calculations based on the work of Stouthamer (54). For the production of 40 g of cells (Y_glycerol_^max^), which contains 20 g of carbon (i.e. 50%), 92 (molecular weight of glycerol) / 36 (molecular mass of carbon in glycerol) × 20 g = 51 g of glycerol was utilized. Consequently, 51 g of glycerol has been assimilated into cell material and 92 – 51 = 41 g of glycerol has been oxidized (dissimilated) to CO_2_. This necessarily assumes that the 51 g of glycerol (55%) assimilated undergo no energy yielding transformation before biosynthetic usage and no other carbon sources were utilized. As the net gain of ATP per glycerol oxidized is 22, the amount of ATP formed from 41 g of glycerol is 41/92 × 22 = 9.8 moles of ATP/mol glycerol utilized. Similar values have been reported for other aerobic bacteria grown in glycerol-limited continuous culture (55, 56). Using these values we calculated the Y_ATP_ values at the respective dilution rates. For example, at a dilution rate of 0.1 h^−1^, the Y_glycerol_ was 34 g [dry weight] of cells/mol glycerol utilized or 34 g [dry weight] of cells/9.8 moles of ATP utilized which is equivalent to a Y_ATP_ value of 3.47 g [dry weight] cells/mol ATP. Based on these calculations, the Y_ATP_ values were in the range 2.90 –3.90 g [dry weight] cells/mol ATP consumed (Figure S6C).

When 1/Y_ATP_ versus 1/dilution rate was plotted the data yielded a straight line (Figure S6D). The intercept on the ordinate was Y_ATP_^max^ corrected for maintenance ATP requirements, and the slope of the function represents the rate of ATP utilization for maintenance functions (mATP). The Y_ATP_^max^ was 3.80 g [dry weight] cells/mol ATP consumed and mATP was 3 mmol of ATP/h/g [dry weight] of cells (Figure S6D). The Y_ATP_^max^ of 3.80 g [dry weight] cells/mol ATP is equivalent to an ATP demand of 263 mmol ATP per gram of newly synthesized biomass. Glucose-limited cultures of *Bacillus subtilis* have a Y_ATP_^max^ of 9.5 g [dry weight] cells/mol ATP, which is equivalent to an ATP demand of 105 mmol per gram or newly synthesized biomass (55). In *E. coli*, the Y_ATP_^max^ value growing in glycerol-limited conditions was 12.7 g cells/mol ATP equivalents(55) which is equal to an ATP demand of 79 mmol ATP per gram or newly synthesized biomass, a value 3.3-fold lower than *M. smegmatis*. The energy requirement for maintenance purposes in *E. coli* was approximately 2 mmol ATP/h/g [dry weight] of cells during energy-limited growth, which is comparable *to M. smegmatis* (3 mmol of ATP/h/g [dry weight] of cells). A Y_ATP_^max^ value of 13.5 (an ATP demand of 74 mmol ATP per gram cells) has been reported for *Bacillus megaterium* grown under the same conditions (56). These comparisons suggest that the synthesis of a mycobacterial cell is an ATP expensive process, which may explain why oxidative phosphorylation by the F_1_F_o_-ATP synthase is essential in mycobacteria and why H_2_ oxidation is primarily used for maintenance rather than to support growth(52).

---

**Figure S1.**
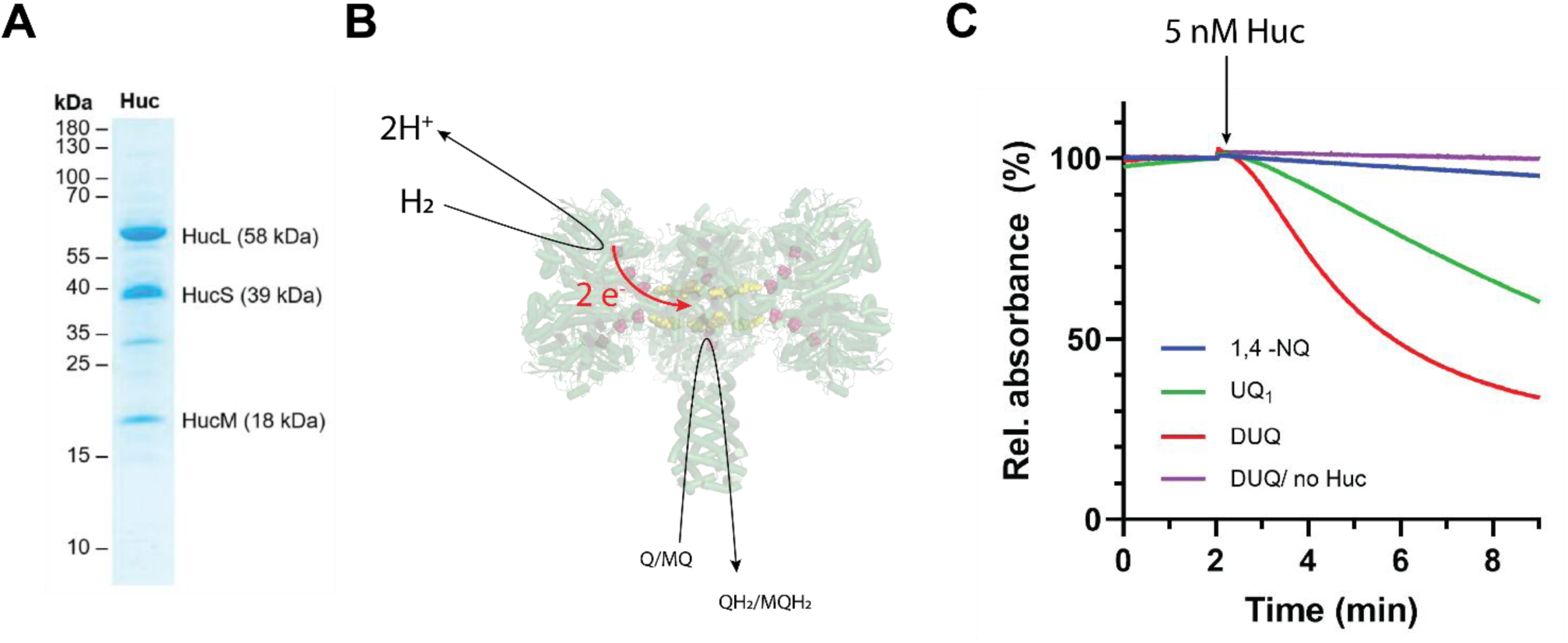
A.) SDS Page of purified Huc B.) Cartoon of quinone reduction experiments in detergent solution under anaerobic conditions. C.) Raw data from Huc-driven quinone reduction measurements, using 50 μM ubiquinoe Q_1_ or decylubiquinone in the presence of ∼500 μM H_2_. Negative control shows no spontaneous reduction of quinone in the absence of H_2_.

**Figure S2.**
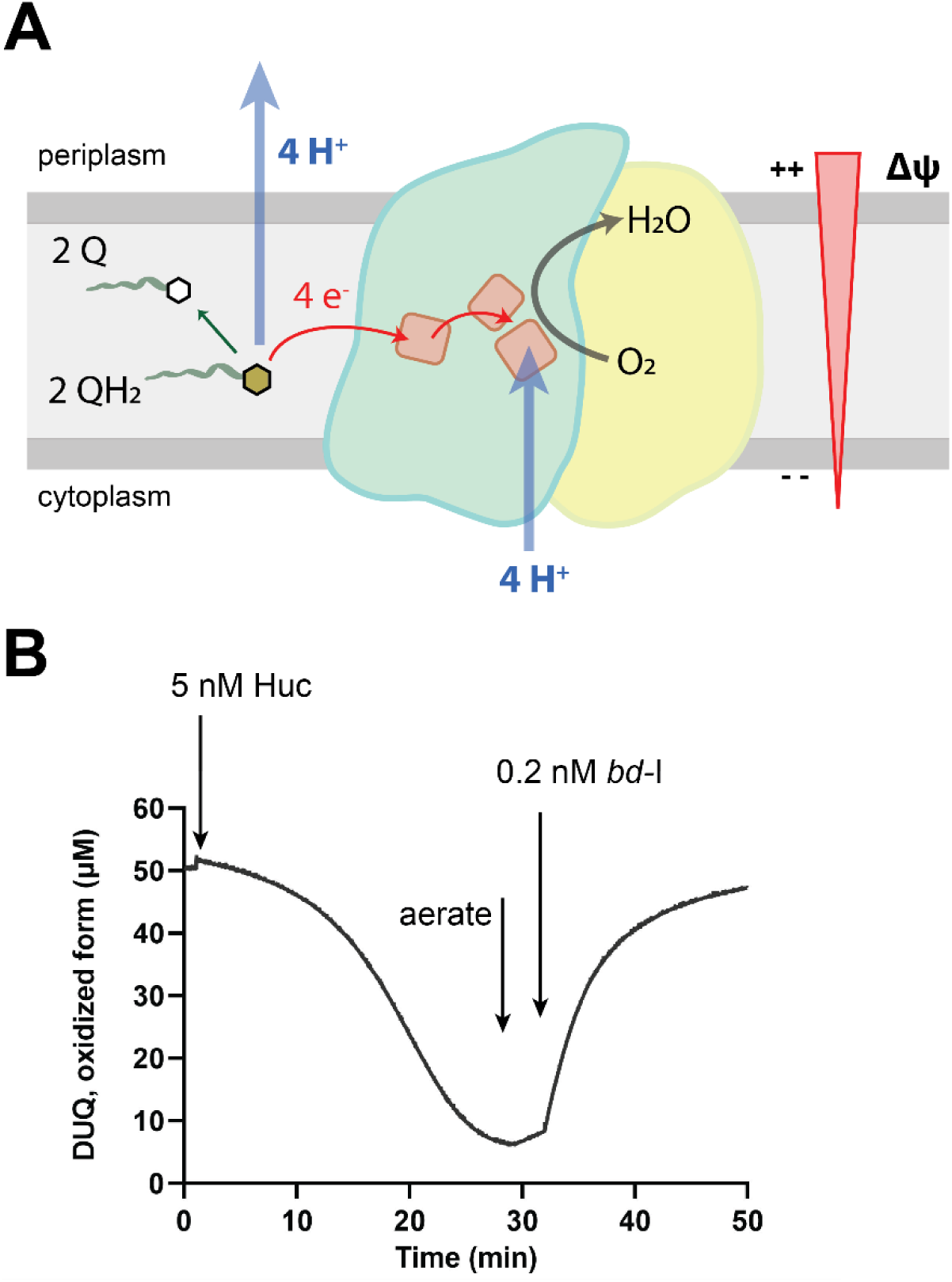
**A.)** Cartoon representation of the energy conserving mechanism of *bd*-I type oxidase. Membrane bound quinones reach the catalytic site in the middle of the membrane and electrons are transferred *via* heme groups to the catalytic site. Protons released from quinol oxidation are released to periplasm in a non-electrogenic process but contributes to scaler proton release. Uptake of protons from the cytoplasm to the catalytic site for water formation is electrogenic and creates a membrane potential Δψ, leading to a net energy conservation of 1H^+^/e^−^. **B.)** Coupling of Huc-driven quinone reduction that is harnessed by *bd*-I oxidase to reduce oxygen to water. Followed is the redox state of 50 μM decylubiquinone at 277 nm. The quinone is reduced under anaerobic condition in the presence of ∼500 μM H_2_. Upon aeration of the cuvette and addition of a 0.2 nM *bd-*I oxidase.

**Figure S3.**
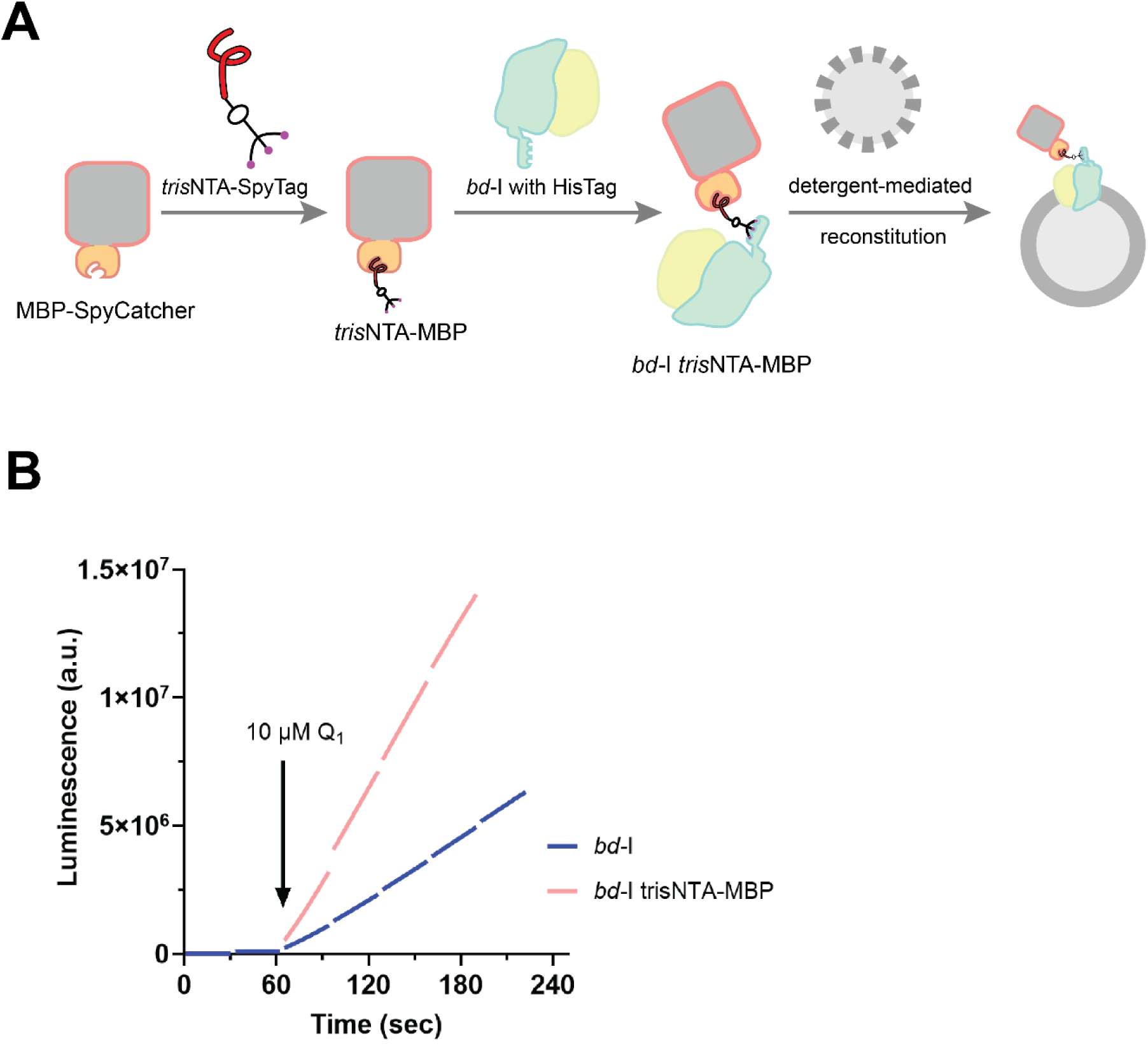
A.) Guided membrane protein reconstitution of *bd*-I oxidase into liposomes. Purifed *bd*-I oxidase is transiently tagged via HisTag with a soluble protein (MBP-SpyCatcher fusion protein that is equipped with *tris*NTA moieties that have a high affinity for HisTag. Reconstitution into liposomes treated with non-solubilizing concentrations of detergent lead to a preferential orientation with the MBP moiety on the outside as described earlier for proteorhodopsin (29). B.) Effect of MBP-moiety on ATP synthesis in liposomes co-reconstituted with ATP synthase, and ATP synthesise was initiated by addition of DTT/UQ_1_ as electron donor and mediator.

**Figure S4.**
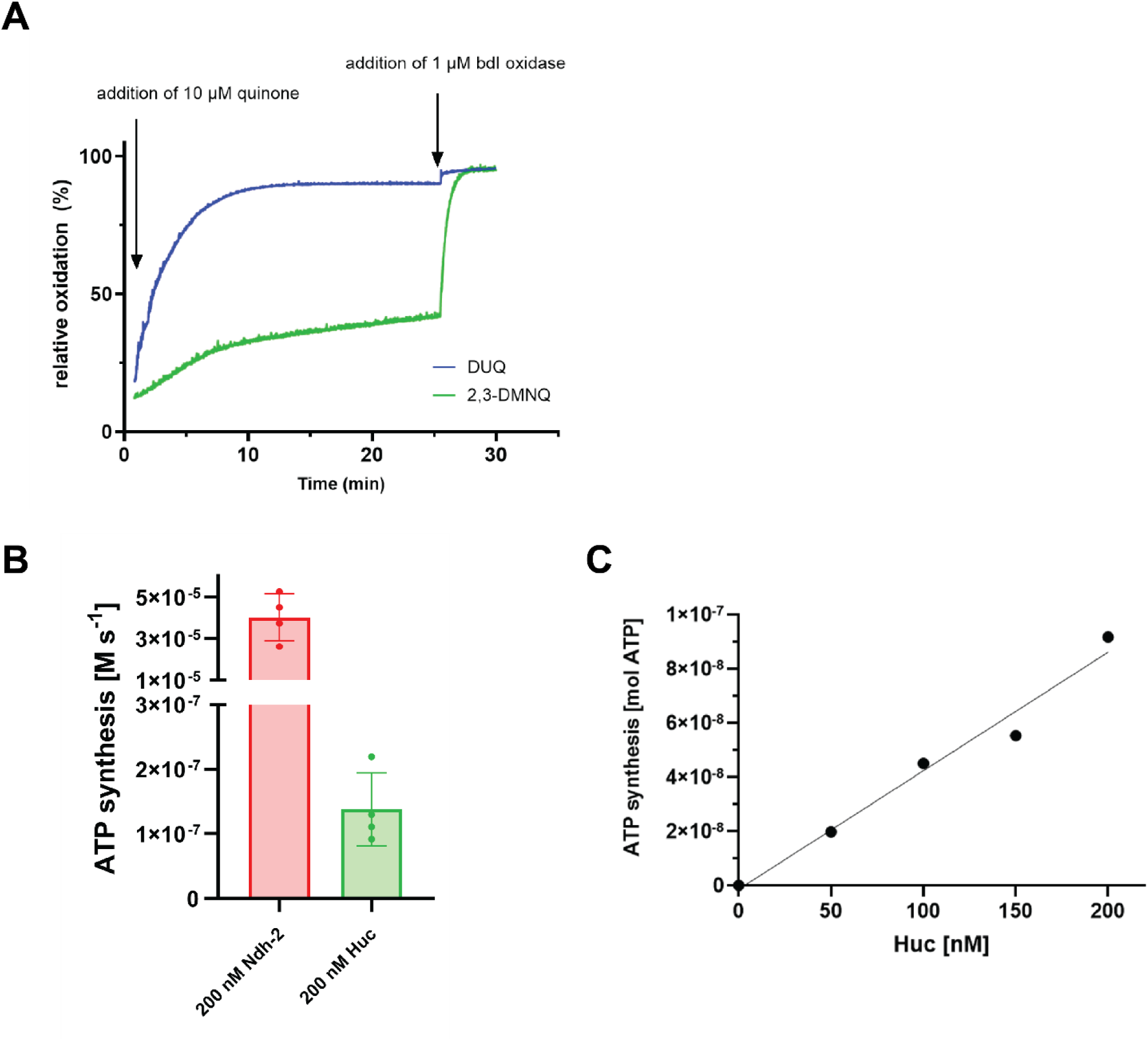
A.) Autooxidation of DUQ and 2,3-DMNQ under aerobic ATP synthesis conditions. B.) Maximal ATP synthesis rates compared for energization by NDH-2 and 1 mM NADH *vs.* Huc/100 μM H_2._ C.) Titration of Huc during ATP synthesis measurements, showing that 200 nM Huc is not sufficient to saturate the liposomes and the Huc reaction is rate limiting.

**Figure S5:**
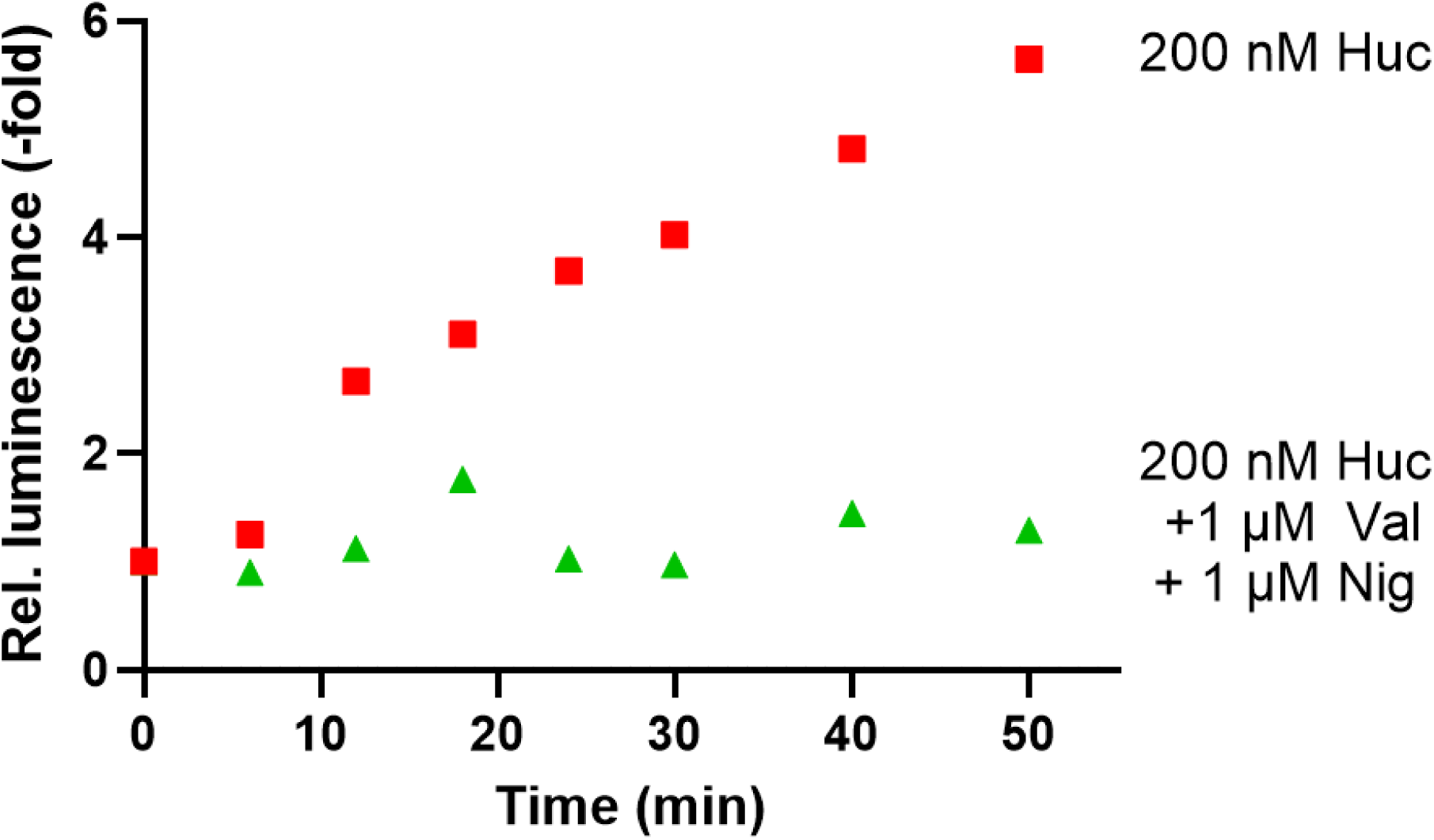
Huc-driven ATP synthesis from air similar to Figure 3A, but with a different Huc preparation. Shown are raw traces from ATP synthesis measurements in an open vessel using 50 μM DMNQ as electron mediator, containing either 200 nM Huc (red), 200 nM Huc and 1 μM valinomycin and 1 μM nigericin (green). Shown are the average values of two technical duplicates.

**Figure S6.**
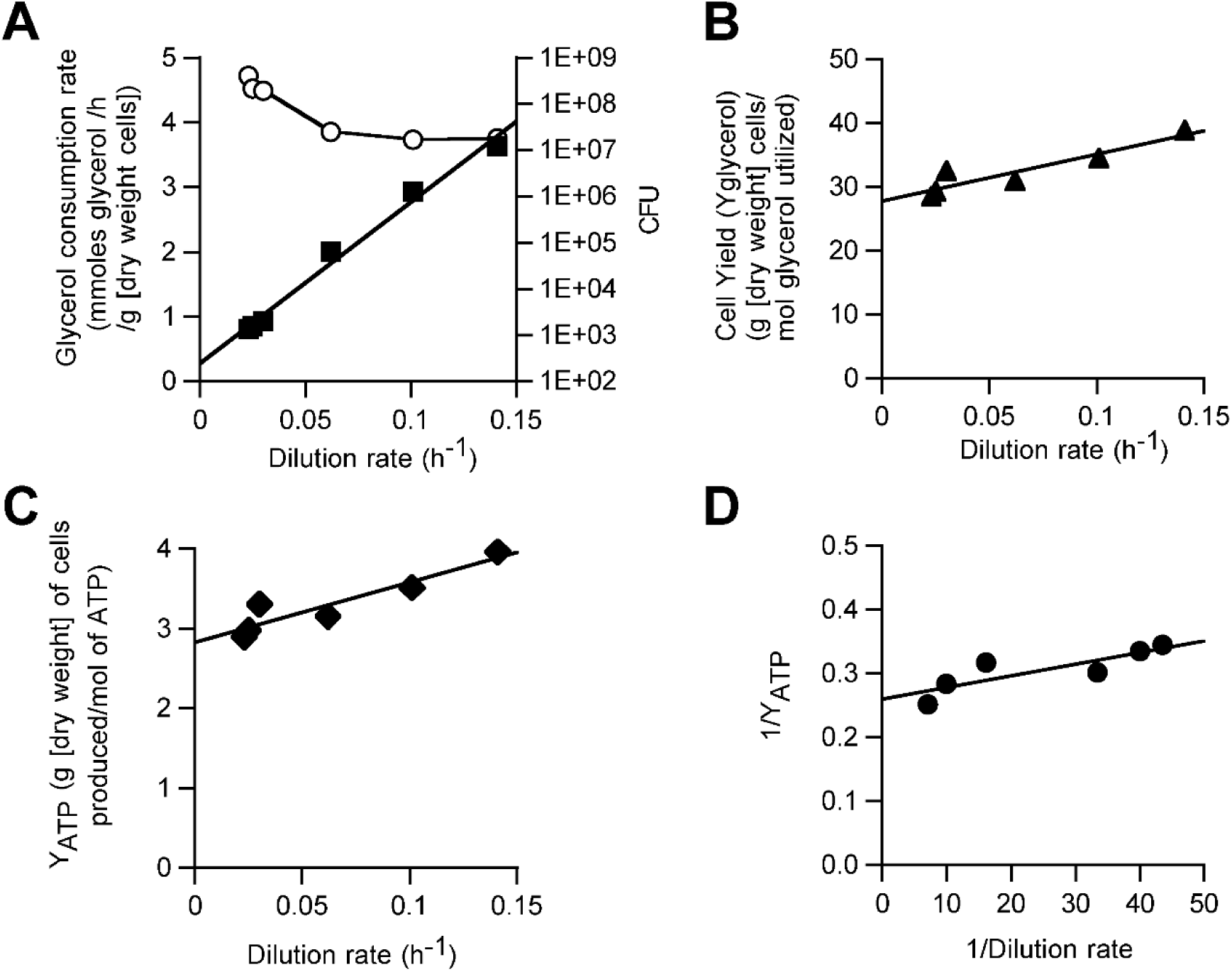
Bioenergetic parameters of *Mycobacterium smegmatis* continuous cultures as a function of dilution rate (controlled by glycerol availability). A) Glycerol consumption rate and cell counts measured by colony forming units (CFU mL^−1^), B) Cell yield per glycerol consumpted (Y_glycerol_), C) Cell yield per ATP consumed (Y_ATP_), D) Maintenance energy requirements (m_ATP_; based on slope of function 1/Y_ATP_ over 1/Dilution rate). The values reported are the means of two to three independent experiments and three technical replicates as each dilution rate and the experimental error associated with the values was less than 20%.

## Notes

### Competing Interest Statement

The authors have declared no competing interest.

## References

1. D. G. Nicholls, S. J. Ferguson, Eds., Bioenergetics - 4th edition (Academic Press, 2013).

2. C. von Ballmoos, A. Wiedenmann, P. Dimroth, Essentials for ATP synthesis by F1F0 ATP synthases. Annu. Rev. Biochem 78, 649–672 (2009).

3. C. Greening, R. Grinter, Microbial oxidation of atmospheric trace gases. Nat Rev Microbiol 20, 513– 528 (2022).

4. S. K. Bay, et al., Microbial aerotrophy enables continuous primary production in diverse cave ecosystems. [Preprint] (2024). Available at: https://www.biorxiv.org/content/10.1101/2024.05.30.596735v1 [Accessed 31 January 2025].

5. A. Kropp, et al., Quinone extraction drives atmospheric carbon monoxide oxidation in bacteria. Nat Chem Biol 1–11 (2025). 10.1038/s41589-025-01836-0.

6. R. Grinter, et al., Structural basis for bacterial energy extraction from atmospheric hydrogen. Nature 615, 541–547 (2023).

7. A. T. Tveit, et al., Widespread soil bacterium that oxidizes atmospheric methane. Proceedings of the National Academy of Sciences 116, 8515–8524 (2019).

8. P. Constant, S. P. Chowdhury, J. Pratscher, R. Conrad, Streptomycetes contributing to atmospheric molecular hydrogen soil uptake are widespread and encode a putative high-affinity [NiFe]-hydrogenase. Environmental Microbiology 12, 821–829 (2010).

9. C. Greening, M. Berney, K. Hards, G. M. Cook, R. Conrad, A soil actinobacterium scavenges atmospheric H2 using two membrane-associated, oxygen-dependent [NiFe] hydrogenases. Proceedings of the National Academy of Sciences 111, 4257–4261 (2014).

10. C. Greening, S. G. Villas-Bôas, J. R. Robson, M. Berney, G. M. Cook, The Growth and Survival of Mycobacterium smegmatis Is Enhanced by Co-Metabolism of Atmospheric H2. PLOS ONE 9, e103034 (2014).

11. Q. Liot, P. Constant, Breathing air to save energy – new insights into the ecophysiological role of high-affinity [NiFe]-hydrogenase in Streptomyces avermitilis. MicrobiologyOpen 5, 47–59 (2016).

12. P. M. Leung, et al., Trace gas oxidation sustains energy needs of a thermophilic archaeon at suboptimal temperatures. Nat Commun 15, 3219 (2024).

13. M. Ji, et al., Atmospheric trace gases support primary production in Antarctic desert surface soil. Nature 552, 400–403 (2017).

14. S. K. Bay, et al., Trace gas oxidizers are widespread and active members of soil microbial communities. Nat Microbiol 6, 246–256 (2021).

15. R. Lappan, et al., Molecular hydrogen in seawater supports growth of diverse marine bacteria. Nat Microbiol 8, 581–595 (2023).

16. D. H. Ehhalt, F. Rohrer, The tropospheric cycle of H2: a critical review. Tellus B: Chemical and Physical Meteorology 61, 500–535 (2009).

17. P. Constant, L. Poissant, R. Villemur, Tropospheric H2 budget and the response of its soil uptake under the changing environment. Science of The Total Environment 407, 1809–1823 (2009).

18. T. Yano, L.-S. Li, E. Weinstein, J.-S. Teh, H. Rubin, Steady-state Kinetics and Inhibitory Action of Antitubercular Phenothiazines on Mycobacterium tuberculosis Type-II NADH-Menaquinone Oxidoreductase (NDH-2) *. Journal of Biological Chemistry 281, 11456–11463 (2006).

19. K. Björklöf, V. Zickermann, M. Finel, Purification of the 45 kDa, membrane bound NADH dehydrogenase of Escherichia coli (NDH-2) and analysis of its interaction with ubiquinone analogues. FEBS Letters 467, 105–110 (2000).

20. W. J. Ingledew, R. K. Poole, The respiratory chains of Escherichia coli. Microbiol Rev 48, 222–71 (1984).

21. P. J. O’Brien, Molecular mechanisms of quinone cytotoxicity. Chemico-Biological Interactions 80, 1–41 (1991).

22. P. Wardman, Reduction Potentials of One-Electron Couples Involving Free Radicals in Aqueous Solution. Journal of Physical and Chemical Reference Data 18, 1637–1755 (1989).

23. A. Theßeling, et al., Homologous bd oxidases share the same architecture but differ in mechanism. Nature communications 10, 5138 (2019).

24. V. B. Borisov, R. B. Gennis, J. Hemp, M. I. Verkhovsky, The cytochrome bd respiratory oxygen reductases. Biochimica et Biophysica Acta (BBA) - Bioenergetics 1807, 1398–1413 (2011).

25. V. B. Borisov, et al., Aerobic respiratory chain of Escherichia coli is not allowed to work in fully uncoupled mode. Proceedings of the National Academy of Sciences 108, 17320–17324 (2011).

26. P. N. Refojo, F. V. Sena, F. Calisto, F. M. Sousa, M. M. Pereira, The plethora of membrane respiratory chains in the phyla of life. Advances in microbial physiology 74, 331–414 (2019).

27. F. Calisto, F. M. Sousa, F. V. Sena, P. N. Refojo, M. M. Pereira, Mechanisms of Energy Transduction by Charge Translocating Membrane Proteins. Chemical reviews 121, 1804–1844 (2021).

28. A. Wiedenmann, P. Dimroth, C. von Ballmoos, Δψ and ΔpH are equivalent driving forces for proton transport through isolated F0 complexes of ATP synthases. Biochimica et Biophysica Acta (BBA) - Bioenergetics 1777, 1301–1310 (2008).

29. A. M. Amati, et al., Overcoming Protein Orientation Mismatch Enables Efficient Nanoscale Light-Driven ATP Production. ACS Synth. Biol. (2024). 10.1021/acssynbio.4c00058.

30. C. von Ballmoos, O. Biner, T. Nilsson, P. Brzezinski, Mimicking respiratory phosphorylation using purified enzymes. Biochim Biophys Acta 1857, 321–31 (2016).

31. S. Deutschmann, et al., Modulating Liposome Surface Charge for Maximized ATP Regeneration in Synthetic Nanovesicles. ACS Synth. Biol. 13, 4061–4073 (2024).

32. B. Wiseman, et al., Structure of a functional obligate complex III(2)IV(2) respiratory supercomplex from Mycobacterium smegmatis. Nature structural & molecular biology 25, 1128–1136 (2018).

33. P. R. F. Cordero, et al., Two uptake hydrogenases differentially interact with the aerobic respiratory chain during mycobacterial growth and persistence. Journal of Biological Chemistry 294, 18980– 18991 (2019).

34. D. Pogoryelov, et al., Engineering rotor ring stoichiometries in the ATP synthase. Proc Natl Acad Sci U S A 109, E1599–608 (2012).

35. L. Preiss, et al., Structure of the mycobacterial ATP synthase Fo rotor ring in complex with the anti-TB drug bedaquiline. Sci Adv 1, e1500106 (2015).

36. A. Meyrat, C. von Ballmoos, ATP synthesis at physiological nucleotide concentrations. Sci Rep 9, 3070 (2019).

37. M. Berney, G. M. Cook, Unique Flexibility in Energy Metabolism Allows Mycobacteria to Combat Starvation and Hypoxia. PLOS ONE 5, e8614 (2010).

38. C. Greening, et al., Persistence of the dominant soil phylum Acidobacteria by trace gas scavenging. Proceedings of the National Academy of Sciences 112, 10497–10502 (2015).

39. Z. F. Islam, et al., Two Chloroflexi classes independently evolved the ability to persist on atmospheric hydrogen and carbon monoxide. The ISME Journal 13, 1801–1813 (2019).

40. L. K. Meredith, et al., Consumption of atmospheric hydrogen during the life cycle of soil-dwelling actinobacteria. Environmental Microbiology Reports 6, 226–238 (2014).

41. M. Berney, C. Greening, K. Hards, D. Collins, G. M. Cook, Three different [] hydrogenases confer metabolic flexibility in the obligate aerobe ycobacterium smegmatis. Environmental Microbiology 16, 318–330 (2014).

42. T. Schmider, et al., Physiological basis for atmospheric methane oxidation and methanotrophic growth on air. Nat Commun 15, 4151 (2024).

43. D. H. Ehhalt, F. Rohrer, The tropospheric cycle of H2: a critical review. Tellus B: Chemical and Physical Meteorology 61, 500–535 (2009).

44. A. M. Amati, S. Graf, S. Deutschmann, N. Dolder, C. von Ballmoos, Current problems and future avenues in proteoliposome research. Biochem Soc Trans 48, 1473–1492 (2020).

45. E. Bailoni, et al., Minimal Out-of-Equilibrium Metabolism for Synthetic Cells: A Membrane Perspective. ACS Synth Biol 12, 922–946 (2023).

46. L. Otrin, et al., Artificial Organelles for Energy Regeneration. Advanced Biosystems 3, 1800323 (2019).

47. A. Abou-Hamdan, et al., Functional design of bacterial superoxide:quinone oxidoreductase. Biochim Biophys Acta Bioenerg 1863, 148583 (2022).

48. J. Hoeser, S. Hong, G. Gehmann, R. B. Gennis, T. Friedrich, Subunit CydX of Escherichia coli cytochrome bd ubiquinol oxidase is essential for assembly and stability of the di-heme active site. FEBS Lett 588, 1537–1541 (2014).

49. J. Eberhardt, D. Santos-Martins, A. F. Tillack, S. Forli, AutoDock Vina 1.2.0: New Docking Methods, Expanded Force Field, and Python Bindings. J. Chem. Inf. Model. 61, 3891–3898 (2021).

50. O. Trott, A. J. Olson, AutoDock Vina: improving the speed and accuracy of docking with a new scoring function, efficient optimization and multithreading. J Comput Chem 31, 455–461 (2010).

51. G. M. Cook, The intracellular pH of the thermophilic bacterium Thermoanaerobacter wiegelii during growth and production of fermentation acids. Extremophiles 4, 279–284 (2000).

52. M. Rao, T. L. Streur, F. E. Aldwell, G. M. Cook, Intracellular pH regulation by Mycobacterium smegmatis and Mycobacterium bovis BCG. Microbiology (Reading) 147, 1017–1024 (2001).

53. M. Daffé, H. Marrakchi, Unraveling the Structure of the Mycobacterial Envelope. Microbiology Spectrum 7, 10.1128/microbiolspec.gpp3-0027–2018 (2019).

54. A. H. Stouthamer, A theoretical study on the amount of ATP required for synthesis of microbial cell material. Antonie Van Leeuwenhoek 39, 545–565 (1973).

55. I. S. Farmer, C. W. Jones, The Energetics of Escherichia coli during Aerobic Growth in Continuous Culture. European Journal of Biochemistry 67, 115–122 (1976).

56. A. J. Downs, C. W. Jones, Energy conservation in Bacillus megaterium. Arch. Microbiol. 105, 159– 167 (1975).

